# Germ-free piglets display variable neuroinflammatory-like perturbations in prefrontal cortical microglia

**DOI:** 10.64898/2026.03.22.713463

**Authors:** Brooke A. Lester, Colin Kelly, Sara N. Henry, Isaac P. Elias, Sophia E. Cevenini, Margaret R. Hendrickson, Taeseo Park, Theodore D. Ashley, Julianna M. Beltz, Julia P. Milner, Alicia M. Pickrell, Paul D. Morton

**Affiliations:** School of Neuroscience, Virginia Polytechnic Institute and State University, Blacksburg, VA 24061 USA; Graduate Program in Translational Biology, Medicine, and Health, Virginia Polytechnic Institute and State University, Roanoke, VA 24016 USA; Department of Pathobiology and Biomedical Sciences, Virginia-Maryland, College of Veterinary Medicine, Virginia Polytechnic Institute and State University, Blacksburg, VA 24061 USA

## Abstract

Communication between gut microbiota and immune cells within the brain is essential for neurotypical development. Specifically, microglia are known to play a key role in regulating and supporting neural progenitor stem cell production during brain development, and are sensitive to changes in the maternal gut microbial composition during perinatal development. Here, we employed a germ-free (GF) porcine paradigm to examine how the absence of the microbiome affects microglial dynamics during a key epoch of brain development. We utilized automated software to evaluate microglial density and morphology across three developmentally significant regions: the ventricular/subventricular zone (VZ/SVZ), the prefrontal subcortical white matter (PFCSWM), and layers II/III of the prefrontal cortex (PFCII-III). We found no significant differences in microglial morphology or density in the VZ/SVZ or PFCSWM. In contrast, the PFCII-III of P16 piglets exhibited an increase in microglia density paired with morphologies indicative of an activated/reactive functional state. Notably, these effects were identified with no overall changes in microglial density in any of the regions assessed. Transcriptomics on RNA isolated from the PFCII-III revealed a significant upregulation of genes related to neuroinflammation, in agreement with a region-specific microglial and immune response in the absence of microbial colonization during postnatal development. Together, these findings build on the limited knowledge available on how microbiota influence brain development in large animal model organisms with high similarities to human brain anatomy and developmental trajectories.

**Significance Statement:** The prefrontal cortex of porcine display unique, ramified microglia which are sensitive to germ-free conditions whereby they display alterations in morphology with a more transcriptionally reactive signature. These findings indicate that microglia are regionally sensitive to stimuli in the periphery, and studies in lissencephalic mammalian models may not be directly correlative to other higher-order species. The neuroanatomical heterogeneity of microglia across species is informative and understudied, but necessary, to draw conclusions on the array of perturbations spanning neurodevelopmental trajectories in health and disease.

## Introduction

The gut microbiome is necessary for healthy development and alterations in gut microbial populations can lead to detrimental effects on metabolic and immune cell function (Borre et al., 2014; Sharon et al., 2016; Lu et al., 2018). The link between the brain and microbiome is an ever-expanding area of research as recent findings suggest the presence and importance of both direct and indirect communication between intestinal microbes, the gut, and the brain (Morais et al., 2021). This communication is supported through microbial colonization that aligns with important landmarks of brain development, including neurogenesis and neuronal migration (Lebovitz et al., 2018; Cowan et al., 2020). Both disruption of this communication and differences in microbial populations and metabolites have been shown to have drastic effects on the brain and are implicated in neurological diseases including autism spectrum disorders (ASD), multiple sclerosis (MS), depression, anxiety, and many neurodegenerative diseases (Lu et al., 2018; Martin et al., 2018; Kim et al., 2019; Chevalier et al., 2020; i and i, 2022; Morton et al., 2023).

Pathologies in the prefrontal cortex contribute to disorders like ASD, depression, and anxiety; however, the role of microbiota during complex cortical development is largely unexplored. In addition, gut dysbiosis is common in preterm birth, representing 10% of all live births in the US, with these patients often displaying reduced cortical volume and suffering from a broad array of neurodevelopmental disorders (Botting et al., 1997; Anderson et al., 2003; Anderson et al., 2004; Dyet et al., 2006; Thompson et al., 2007; Delobel-Ayoub et al., 2009; Ball et al., 2012; de Kieviet et al., 2012; Molnar and Rutherford, 2013). While normal delivery at birth marks the onset of gut colonization with maternal microbes, the microbiota of preterm infants is largely shaped by other factors including antibiotics and the external environment within the neonatal intensive care unit (Brooks et al., 2014). Therefore, a strong understanding of how microbiota or lack thereof influence cortical development is of significance.

As resident macrophages continuously surveil the central nervous system (CNS), microglia have been well studied in the context of their immunological roles in the adult brain. More recently, studies have indicated that microglia also play critical roles in complex neurodevelopmental programs underlying brain development and are key players in the progression of developmental diseases (Cowan and Petri, 2018; Lukens and Eyo, 2022). The human cerebral cortex is largely built by neurons originating in the ventricular zone (VZ) and subventricular zone (SVZ) during pre- and perinatal development; tight regulation of cell production is critical for proper brain development and function (Lehtinen and Walsh, 2011; Lui et al., 2011; Lim and Alvarez-Buylla, 2014; Beattie and Hippenmeyer, 2017). During this defined critical period, microglia colonize these niches and actively phagocytose neural stem progenitor cells (NSPCs) preventing overproduction of newborn neurons, and further provide trophic support to the niche and resident NSPCs (Cunningham et al., 2013; Shigemoto-Mogami et al., 2014; Ribeiro Xavier et al., 2015; Morton et al., 2018). In addition, the gut microbiome globally regulates microglia in a sexually dimorphic manner pre- and postnatally in germ-free (GF) mice (Thion et al., 2018) as well as in a region-specific manner (Paolicelli et al., 2022; Huang et al., 2023). Microglia morphology is extremely dynamic, sensitive to environmental cues, and provides information that imply functional status.

Previous studies found that microglia in adult raised GF mice had increased numbers and morphological complexity than conventionally housed counterparts (Erny et al., 2015); an alteration that was also observed at birth, but not preterm (Castillo-Ruiz et al., 2018). Our study sought to examine similar parameters in the VZ/SVZ of neonatal piglets, where microglia are found in abundance, and prefrontal cortex (PFC). Although region specific differences were clear in microglia density and morphology, our data suggests that microglia display differences and a more activated morphology in GF paradigm in only the prefrontal cortex. These results were validated transcriptomically, with upregulated differentially expressed proinflammatory genes in the PFC indicating a more inflammatory/active phenotype. These results indicate that species specific, regionally different microglia most likely exert divergent effects on neurodevelopmental trajectories and other vertebrate model systems are required to mechanistically tease out the effects of the microbiome and its metabolites on the brain.

## Material and Methods

### Animals

Domestic pigs (*Sus scrofa*, Landrace × Yorkshire crossbreed) including males and females were used for all experiments. A total of 20 piglets were used in this study (P0, n = 1; P16, n = 18, P42, n = 1): histology (Control, n = 4F, 2M; GF, n = 5F, 1M) and RNA isolation (Control, n = 2F, 1M; GF, n = 1F, 2M). Derivation, rearing, and euthanasia of control and GF animals was as previously described (Ahmed et al., 2021). Pigs were received from Oak Hill Genetics, IL, and the Swine Agricultural Research and Extension Center at Virginia Tech. All experiments were conducted in accordance with the NIH Guide for the Care and Use of Laboratory Animals and carried out under the approval of the Virginia Tech Institutional Animal Care and Use Committee.

Mice were housed in a pathogen-free facility on a 12h light/dark cycle at Virginia Tech at a maximum of five per cage and provided standard rodent diet and water *ad libitum*. C57BL/6J WT mixed sex mice were purchased from Jackson Laboratories. All mouse experiments were conducted in accordance with the NIH guide for the Care and Use of Laboratory Animals and under the approval of the Virginia Tech Institutional Animal Care and Use Committee (IACUC).

### Germ-free Paradigm

Control and GF piglets were born from different parents. GF piglets were derived via hysterectomy from near-term sows 1 day prior to term as previously described (Ahmed et al., 2021). To prevent iron deficiencies, 150 mg of iron dextran (Patterson Veterinary Supply) was administered intramuscularly immediately following birth. Piglets were housed in sterile isolators with dimensions (24” × 42”× 24”) designed to accommodate up to four piglets. The piglets were able to hear and smell other animals housed in the same isolator and one toy was provided per chamber. Animals were kept on a 12:12 h light: dark cycle. The room temperature is maintained between 93 and 95°F initially and then decreased 2°F/week for the remainder of the study. Pigs were fed a sterile commercial diet (Hershey’s UHT milk) three times a day; through individual feeders provided for each animal to ensure a controlled diet that increased in volume to accommodate growth. Piglets were closely monitored throughout the day for any clinical signs of concern by the Teaching and Research Animal Care Support Unit (TRACSS) and/or an on-site/call veterinarian. Animals were weighed weekly, and GF status was confirmed biweekly by sterility tests of rectal swabs for aerobic and anaerobic microbes using blood agar and thioglycolate media, respectively; additionally, the isolators were swabbed and tested in the same manner. All items (e.g., weighing instruments, swabs, toys, etc.) were sterilized with Spor-Klenz® prior to introduction to the isolators. Healthy, GF animals were euthanized in accordance with AVMA Guidelines for the Euthanasia of Animals for the following experimental procedures. Control piglets were delivered vaginally and raised in a conventional agricultural environment. P0 piglets were euthanized shortly following birth via hysterectomy.

### Euthanasia

Animals were euthanized by transcardial perfusion. Animals were first sedated. Pre-anesthesia was administered, and anesthesia was provided to prevent pain during the procedure. An absence of interdigital pain reflex was verified before beginning the procedure. The thoracic cavity was opened to access the heart, and a perfusion cannula was inserted into the aorta through the left ventricle, and an incision was then made in the right auricle.

### Tissue Preparation and Immunohistochemistry

Quickly following euthanasia, brains were removed from the skull, weighed, and cut at 5 mm intervals on a custom-designed brain mold, and fixed at 4°C in 4% paraformaldehyde (0.1 M PBS) for 72 h. After fixation, tissue slabs were cryoprotected at 4°C in a sucrose gradient of 20% and 30% (0.1 M PBS) until sunken. Tissue was embedded in OCT compound (Tissue-Plus #4585) and stored at −80°C for subsequent serial sectioning on a Thermo Scientific CryoStar NX50 cryostat. Coronal tissue sections (50 μm) were collected for immunohistochemical procedures.

For immunohistochemistry, sections were incubated with PBST (PBS +.1% Triton X-100) for 20 minutes to permeabilize the tissue. Sections were blocked with 5% donkey serum, 1% BSA in PBST for 1 hour, and then stained with Iba1 (Wako, catalog #NC9288364), Ki67 (Abcam, catalog #ab16667), and CD38 (Proteintech, catalog #60006-1-Ig) overnight at 4°C. Conjugated Alexafluor secondary antibodies (Jackson ImmunoResearch) diluted in blocking buffer were incubated on sections for 1 hour at room temperature. After a final PBS wash, the sections were counterstained with DAPI (Sigma) and cover-slipped using fluorescence anti-fade mounting medium (Southern Biotech).

### Image Acquisition

Following immunostaining, high resolution Z-stack images (25 µm, 0.5 µm step size) were acquired using a Nikon C2 confocal laser scanning microscope (Nikon Instruments, 545 Melville, NY) at 20× or 40× magnification. The Nikon LUN4 has a four-line solid-state laser system with Perfect Focus and DU3 High Sensitivity Detector System. The CFI60 Apochromat Lambda S 40× water immersion objective lens, N.A. 1.25, W.D. 0.16–0.20 mm, F.O.V. 22 mm, DIC, correction collar 0.15–0.19 mm, spring-loaded and a Plan Apto 20x air objective, N.A. 0.75, W.D. 1.00 mm were used for confocal image acquisition. Nikon NIS-Element Package and ImageJ were also used for image analysis. Contours of the SVZ, VZ, PFCII-III, and PFCSWM were drawn by a single unbiased experimenter using a reference atlas. Images were then compressed into a single channel tif file to work with the Microglia Morphology software.

### Microglial Morphology Analysis

For morphological analysis of microglia, images were input into a high throughput analysis pipeline (Kim et al., 2024). A 40× magnification z-stack image was used to assess Iba1^+^ microglia reactivity. The maximum intensity projection image was then loaded into ImageJ, where individual microglial morphology states were distinguished using the open-source macro tool MicrogliaMorphology35. Analysis was performed per the tool’s GitHub protocol. Briefly, the MicrogliaMorphology_BioVoxxel plugin was used to determine thresholding parameters for each image. The lowest intensity point of three random microglia were manually selected to properly determine which thresholding set was best suited for further image analysis. After thresholding, the MicrogliaMorphology_Program plugin was used to determine gating for the single-cell area. Five particles in the image that were considered too small to be single cells were manually selected, followed by five particles that were considered too big, such as overlapping cells. From here, the macro was fully automated and outputs single-cell images, skeleton analysis, and fractal analysis via FracLac. The final outputs from both MicrogliaMorphology and FracLac were merged in RStudio with the MicrogliaMorphologyR package to output a .csv file containing all 27 morphological features for each cell. Using the MicrogliaMorphologyR package, clusters were determined by PCA analysis followed by k-means clustering. The ColorByCluster function mapped each cluster as a different color, which was output onto the original image for visual assessment of cluster identities. For more detailed information on the MicrogliaMorphology macro and protocol, see https://github.com/ciernialab/MicrogliaMorphology.

### RNA Sequencing

For gray matter tissue, animals were deeply anesthetized with a ketamine (500 mg/kg)/xylazine (10 mg/kg) cocktail and hand perfused with cold PBS to remove blood. Tissue was rapidly dissected in ice-cold PBS and stored in RNAlater (Sigma-Aldrich) until RNA extraction. RNA was isolated with the RNeasy Mini Kit (Qiagen) per the manufacturer’s protocol. Isolated RNA was sent to MedGenome (MedGenome Inc.) for RNA sequencing using the Illumina TrueSeq stranded mRNA kit for library preparation and the NovaSeq system for sequencing. RNA-seq data has been deposited on GEO depository: GSE306007.

Bases with quality scores less than 30 and adapters were trimmed from raw sequencing reads by Trim Galore (v0.6.4). After trimming, only reads with length greater than 30 bp were mapped to mm10 by STAR (v2.7.3a). Raw counts for each gene were output by HTSeq (v0.11.2). Raw counts were normalized and used to identify differentially expressed genes (DEGs) by DESeq2 (v1.42.1) in R Studio.

Genes were filtered by a p-value less than 0.05 and at least a 1.2-fold change to be considered differentially expressed. For cortical tissue samples, genes were filtered by an adjusted p-value less than 0.05 and at least a 1.5-fold change. DEGs from gray matter, determined using the thresholds above, were submitted through Ingenuity Pathway Analysis (Qiagen) software for analyses of upstream regulators and top biological functions. The top 10 inhibited and top 10 activated upstream regulators were used to generate the associated bar plot. Top Biological Functions were determined and selected from the Diseases and Functions output, and visually reconstructed in BioRender.

### Statistical Analysis

Appropriate sample size was determined using ANOVA power analysis for 4 groups suggesting 6 animals per group would provide a power of 0.80 at a 0.05 alpha and effect size d=.9. Two-way analysis of variance (ANOVA) was done for groups and regions (two or more), whereas a two-tailed unpaired Student’s t-test was performed for individual comparisons. Significance was determined if p-value was <0.05. Data were expressed in bar graphs as the mean ± standard error of mean (SEM). Dots are representative of the analysis of an individual animal. Significance was shown as *p ≤ 0.05, **p ≤ 0.01, ***p ≤ 0.001. All statistical tests are reported in the figure legends.

## Results

### Microglia are abundant and proximal to mitotic stem/progenitor cells in the postnatal swine SVZ

In rodents, it is well established that microglia begin to exit the SVZ following the peak wave (∼E10) of neurogenesis and decline in numbers within this neurogenic niche as they continue to seed distal domains including subcortical white matter tracts and cortical grey matter (Shigemoto-Mogami et al., 2014); however almost nothing is known regarding the presence or activation status of microglia in the postnatal swine SVZ. Microglial numbers, distribution, and morphology within the SVZ domain (anterior SVZ) known to supply the swine prefrontal cortex with postnatally generated neurons, were evaluated by immunohistochemistry to visualize microglia (Iba1^+^) in relation to mitotic (Ki67^+^) stem/progenitor cells within the SVZ at birth and the age equivalence of toddlerhood in mice and piglets (Workman et al., 2013; Sutkus et al., 2025).

In agreeance with previous reports, the mouse SVZ remained highly mitotic on the day of birth (P0) and into early toddlerhood (P30) with few microglia present indicating microglial emigration to parenchymal tissue *in utero* (**Fig. 1A**). Within the piglet SVZ, microglia were abundant at P0 with a notable decline in numbers by P42 when they were in similar abundance within the surrounding parenchyma suggesting a protracted postnatal exodus of microglia from the SVZ (**Fig. 1B**). Within both species, microglia were endowed with ameboid-like morphologies at birth possibly indicating an activated or immature state; this persisted in the mouse parenchymal tissue surrounding the SVZ during toddlerhood (**Fig. 1A,B**). However, microglia appeared to show higher morphological complexity in both domains assessed in the porcine equivalent (**Fig. 1A,B**).

**Figure 1.**
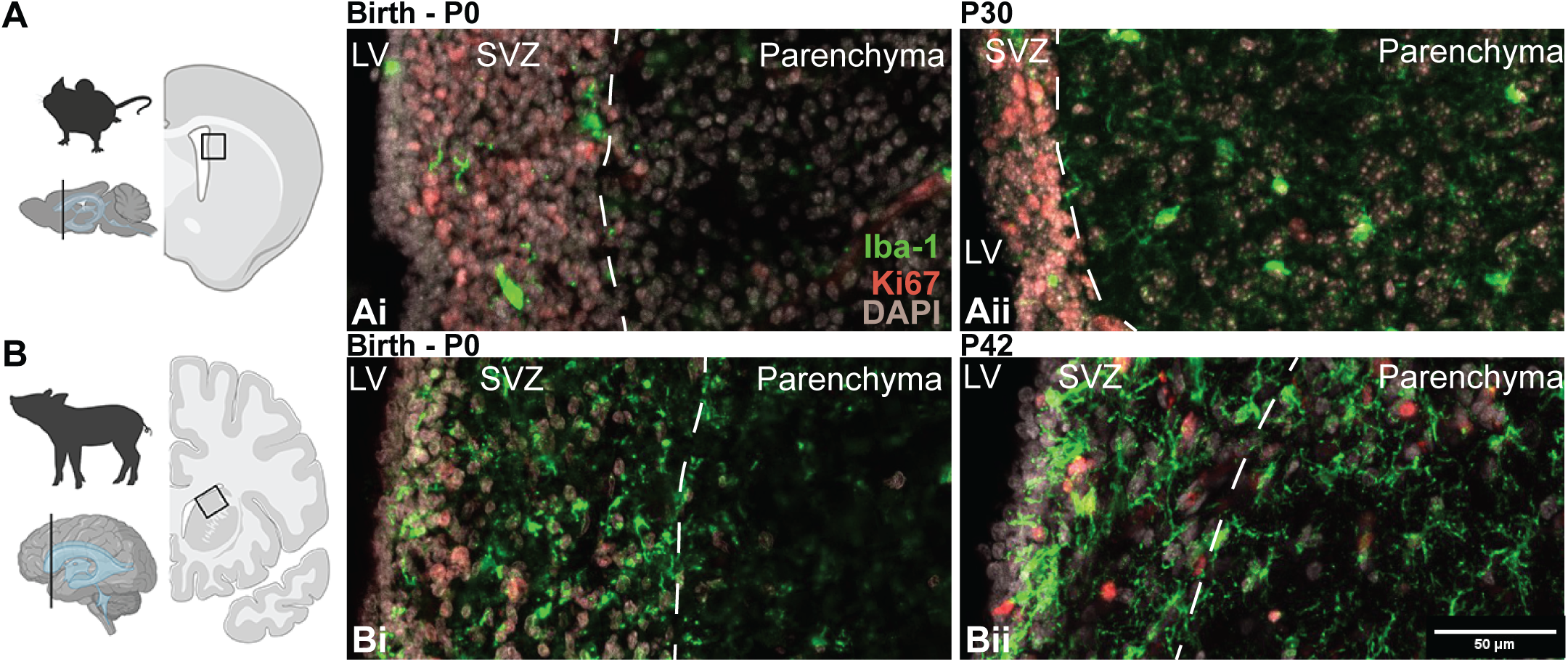
Microglia are abundant and proximal to mitotic stem/progenitor cells in the postnatal swine SVZ. (**A-B**) Immunostains of the P0 and P30 mouse **(A)** and P0 and P42 piglet **(B)** SVZ on the coronal plane of the frontal cortex, illustrating Iba1^+^ microglia (green), Ki67^+^ proliferating cells (red), and DAPI^+^ nuclei (grey). Scale bar = 50 um. Images captured at 20X. Biorender cartoon illustrates species, brain, coronal plane compared (vertical line), and region of SVZ captured (black boxed areas). White lines in z-projections mark the SVZ boundaries.

Together these data illustrate stark differences in microglial residency and morphologies between species which merits further investigation into whether there is overlap between the antenatal role microglia play in buffering stem/progenitor cell numbers and their postnatal roles in immune surveillance and synaptic pruning in higher order species where neurogenesis is robustly ongoing following birth.

### Regional heterogeneity in microglial numbers during neurotypical swine brain development

Differences in the number of microglia - along with distinct morphological features indicative of function - have been documented across various neuroanatomical domains throughout epochs of rodent brain development (De Biase et al., 2017; Hammond et al., 2019). Following birth, the porcine and human SVZ remain neurogenic as newborn interneurons continue to migrate to the prefrontal cortex (Paredes et al., 2016; Morton et al., 2017). Similar to humans, myelination of the porcine brain begins antenatally with a mixed population of oligodendrocyte progenitors and their mature progeny present at birth (Sobierajski et al., 2023). Considering these phylogenetically conserved similarities, we focused on three distinct regions of the neonatal swine brain at P16: (i) ventricular/subventricular zone (VZ/SVZ), (ii) prefrontal subcortical white matter (PFCSWM), and (iii) the upper layers of the prefrontal cortex (PFCII-III).

An unbiased approach utilizing Microglial Morphology software (Kim et al., 2024) was employed to automate evaluation of microglial morphology and density on immunolabeled coronal sections from healthy P16 brains **(Fig. 2**). Quantification of the number of Iba1^+^ microglia relative to the volume of each region of interest, revealed significantly fewer microglia residing in the PFC when compared to the VZ/SVZ and PFCSWM (*F*_(2,13)_ = 9.45, *p* = 0.0029, one-way ANOVA) (**Fig. 2A**). When comparing the numbers of microglia with a “ramified” morphology associated with a resting/homeostatic state, we found no significant differences across regions (*F*_(2,13)_ = 3.35, *p* = 0.067, one-way ANOVA) (**Fig. 2B**). Next, “activated” microglia were classified as the sum of microglia displaying non-ramified features including rod-like, ameboid, or hypertrophic. We found a significant reduction in activated microglia in the PFC compared with the VZ/SVZ and PFCSWM (*F*_(2,13)_ = 44.80, *p* = 0.0000015, one-way ANOVA) (**Fig. 2C**) Regional representative P16 immunostains for each region can be seen in **Figure 2D**.

**Figure 2.**
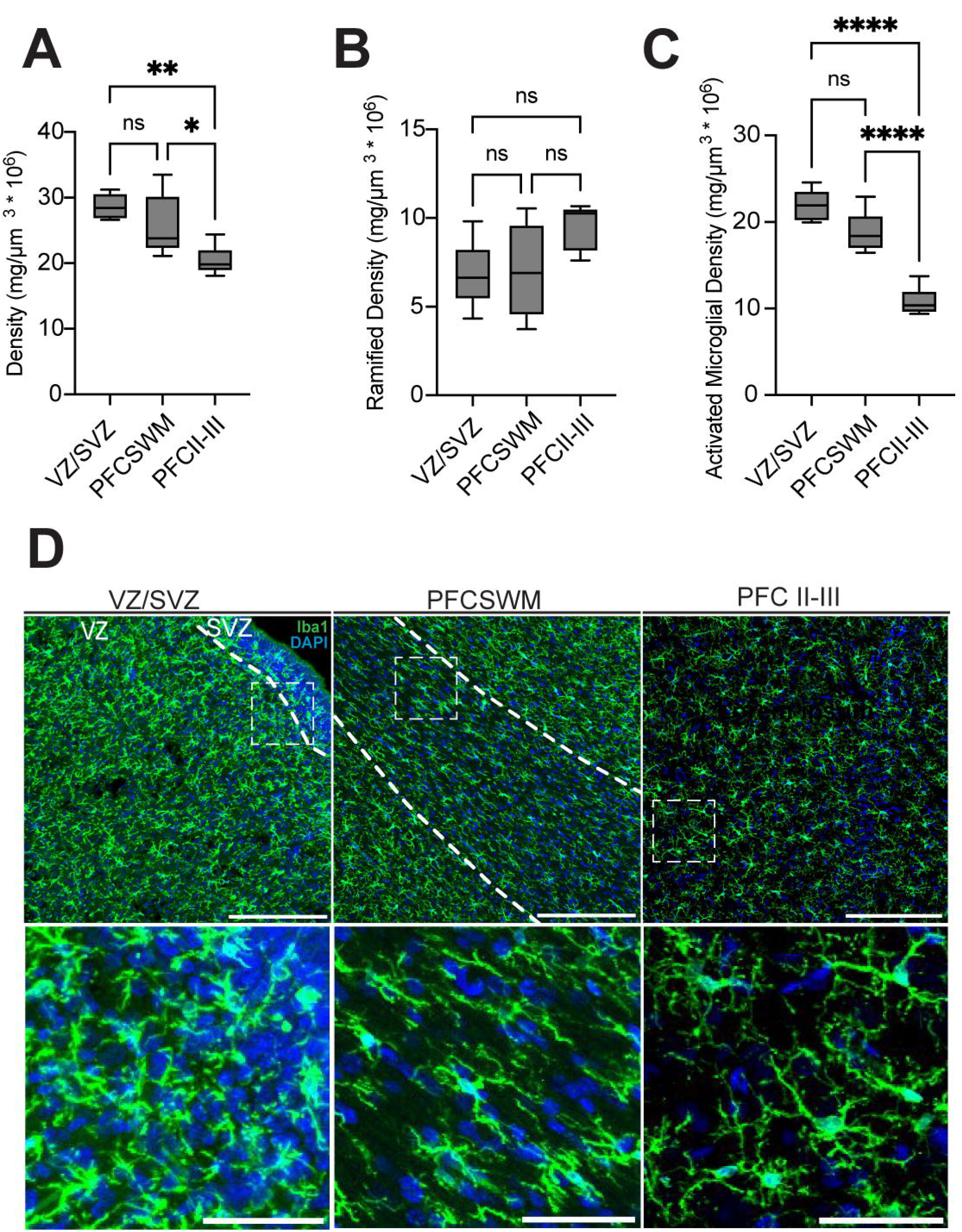
Germ-free conditions have no significant impact on microglial numbers in early postnatal brain development. **(A-C)** Comparisons of microglial density between VZ/SVZ, PFCSWM, and PFCII-III in control tissue overall **(A),** when looking at “resting” (ramified) microglia **(B)**, and when looking at “activated” microglia **(C)**, gathered by MicrogliaMorphology software (density calculated as microglia/µm^3^*10^6^). **(D)** Immunostains of the VZ/SVZ, PFCSWM, and PFCII-III in P16 control piglets. White dashed lines mark regional boundaries. Scale bars, 200µm. Insets 50µm. Error bars indicate mean ± SEM. ns *=* non-significant (*p* > 0.05), \**p* < 0.05, \*\**p* < 0.01, \*\*\**p <* 0.001, \*\*\*\**p* < 0.0001.

Together these data indicate a gradient in microglial cell numbers with an abundance near the lateral ventricles where they initially seeded the brain and scarcer numbers as they migrate long distances to populate the prefrontal cortex. These regional differences are accounted for by a subpopulation of activated microglia which includes not only morphological features of immune cell activation and phagocytosis but also microglial cell migration. Considering that (i) birth provides an immune challenge as the newborn is exposed to maternal microbes, (ii) microglia are sensitive to gut dysbiosis, and (iii) display regional heterogeneity amongst the activated class during neurotypical development, we reasoned that microglial dynamics would be disrupted in germ-free piglets.

### Germ-free conditions have no significant impact on microglial numbers in early postnatal brain development

We previously employed a germ-free (GF) swine paradigm to determine the impact of microbiota on white matter development during early postnatal development (P16) (Ahmed et al., 2021). Utilizing the same GF paradigm, we next assessed microglial cell density within our three regions of interest (**Extended Data Fig. 2-1**). Quantification demonstrated no significant differences within the VZ/SVZ, PFCSWM, or PFCII-III when compared to conventional controls (*t*_(9)_ *=* 1.274, *p* = 0.2345, unpaired *t* test), (*t*_(10)_ *=* 0.9853, *p* = 0.3477, unpaired *t* test), (*t*_(10)_ *=* 2.042, *p* = 0.0684, unpaired *t* test) (**Extended Data Fig. 2-1 A-C**). These findings indicate that the microbiome has no influence over microglial production or survival within each region of interest assessed during early postnatal brain development. In addition, our data suggest no hindrance in the migratory capacity of microglia from the VZ/SVZ to the cortex in the absence of microbial colonization. With no significant changes in microglial population density, we next aimed to assess functional alterations within each key region of interest.

### GF paradigm does not affect microglial morphology in the VZ/SVZ

During early stages of embryonic brain development, microglia enter the brain via the ventricles, play a critical role in buffering stem cell production within the neighboring VZ/SVZ, and subsequently migrate to seed the brain in an inside out manner. Emigration from the VZ/SVZ is largely complete at birth in mice (**Fig. 1**); however, microglia remain within the VZ/SVZ for several weeks (**Fig. 1**) following parturition in piglets. In addition, the anterior dorsolateral region of the P16 swine VZ/SVZ is a highly neurogenic niche, like their human age equivalent, that provides newborn interneurons to the upper layers of the developing prefrontal cortex. The VZ/SVZ is a highly vascularized region of the brain providing a microenvironment enriched in both blood and cerebral spinal fluid-derived circulating factors. Therefore, we reasoned there may be alterations in microglial morphology within this region in GF conditions that could impact multiple stages of cortical development.

Quantification of several output measures generated by Microglial Morphology software revealed no significant differences between control and GF piglets within the VZ/SVZ at P16 (**Fig. 3**). Overall, microglial density between control and GF animals were not significantly different when grouped as “resting” or “activated” (activation state□×□treatment, *F*_(1,16)_□=□0.078, *p□*=□0.783; activation state, *F*_(1,16)_□=□150.1, *p□*=□0.0000000015; treatment, *F*_(1,16)_□=□1.415, *p□*=□0.252, two-way ANOVA), (morphological state□×□treatment, *F*_(3,36)_□=□0.3561, *p□*=□0.785; morphological state, *F*_(3,36)_□=□6.446, *p* = 0.0013; treatment, *F*_(1,36)_□=□1.555, *p□*=□0.2205, two-way ANOVA), (*t*_(9)_ = 1.093, *p* = 0.3026, unpaired *t* test) (**Fig. 3A-C)**. Further, no differences were found between average branch length, branch number, or cell area (*t*_(10)_ *=* 0.00011, *p* = 0.999, unpaired *t* test), (*t*_(10)_ *=* 0.046, *p* = 0.965, unpaired *t* test), (*t*_(10)_ *=* 0.0999, *p* = 0.922, unpaired *t* test) (**Fig. 3D-F**). Representative P16 immunostain for VZ/SVZ can be seen in **Figure 3G**. Taken together, these results demonstrate that GF conditions have no effect on microglial morphology in the VZ/SVZ.

**Figure 3.**
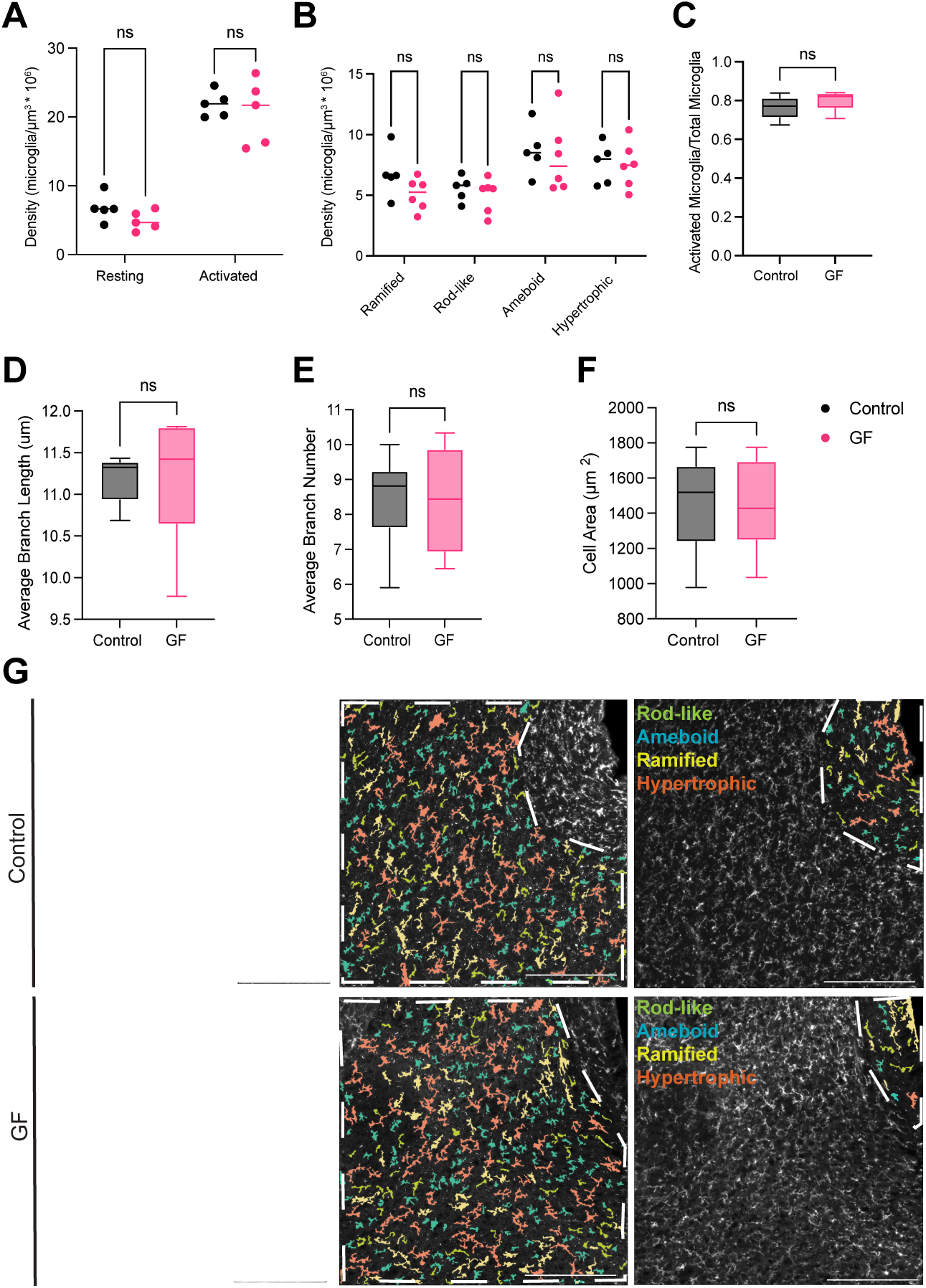
GF paradigm does not affect microglial morphology in the VZ/SVZ. Analyses using microglial morphology data assessed by MicrogliaMorphology software program within the VZ/SVZ. **A,** Quantified density of activated and resting microglia between control (black) and GF (pink) tissue. **B,** Quantified control and GF microglial density within morphological groupings (ramified, rod-like, hypertrophic, ameboid). **C,** Proportion of activated microglia out of total microglia between control and GF tissue. Quantified average branch length (µm)**(D)**, branch number **(E)**, and cell area (µm^3^)**(F)** of individuals between Control and GF groups. **G,** Immunohistochemical images of the P16 VZ/SVZ (first column), the SVZ (second column), and the VZ (third column), in control (upper row) and GF (lower row) tissue. Individual microglia labeled by MicrogliaMorphology Colorbycluster macro as rod-like (green), ameboid (blue), ramified (yellow), or hypertrophic (pink). White dashed lines indicate regional boundaries. Scale bars, 200µm. Density is calculated as microglia/µm^3^*10^6^. Error bars indicate mean ± SEM. ns *=* non-significant (*p* > 0.05).

### Microglia display a significant reduction in branch length in the PFCSWM under germ-free conditions

We previously demonstrated a significant reduction in the numbers of proliferating oligodendrocyte progenitor cells (OPCs) within the subcortical white matter tracts of the prefrontal cortex (PFCSWM) in GF piglets at 16 days of age (Ahmed et al., 2021). Recent evidence suggests that microglia may play a key role in phagocytosing OPCs within the corpus callosum during the early stages of postnatal myelination in mice (Irfan et al., 2022); purportedly to prevent overproduction of oligodendrocytes, thus preserving a balanced ratio to the multiple axons they each myelinate. Therefore, we reasoned that microglia may actively over prune OPCs within the PFCSWM of the developing GF brain and would display morphological differences skewed towards a state of activation.

In the PFCSWM (**Fig. 4**), microglial density between control and GF animals were not significantly different when grouped as “resting” or “activated” (activation state□×□treatment, *F*_(1,16)_□=□0.756, *p□*=□0.397; activation state, *F*_(1,16)_□=□112.8, *p□*=□0.0000000117; treatment, *F*_(1,16)_□=□2.941, *p□*=□0.106, two-way ANOVA) (**Fig. 4A**), grouped by morphology (morphological state□×□treatment, *F*_(3,32)_□=□3.022, *p□*=□0.044; morphological state, *F*_(3,32)□_=□9.889, *p□*=□0.0000913; treatment, *F*_(1,32)_□=□2.584, *p□*=□0.118, two-way ANOVA) (**Fig. 4B**), or in their proportion of activated microglia (*t*_(8)_ = 2.011, *p* = 0.0792, unpaired *t* test) **(Fig. 4C)**. We found a significant reduction in the average branch length of microglia within the PFCSWM of GF piglets (*t*_(8)_ = 2.727, *p* = 0.026, unpaired *t* test) (**Fig. 4D**), which is often regarded as a feature of activated microglia and inflammatory cytokine release (Madry et al., 2018). Lastly, we found no significant differences between control and GF tissues in average number of branches (*t*_(8)_ *=* 1.346, *p* = 0.2153, unpaired *t* test) (**Fig. 4E**) or average cell area (*t*_(8)_ *=* 1.140, *p* = 0.2871, unpaired *t* test) (**Fig. 4F**) of Iba1^+^ microglia. A representative P16 immunostain for PFCSWM can be seen in **Figure 4G**. Overall, a significant reduction in branch length (**Fig. 4D**) paired with no changes in microglia cell density (**Fig. 2E**) suggests that microglia are produced at normal rates and seed the PFCSWM tracts with neurotypical numbers but are in a more functionally active state that may be indicative of protracted phagocytosis of local OPC pools.

**Figure 4.**
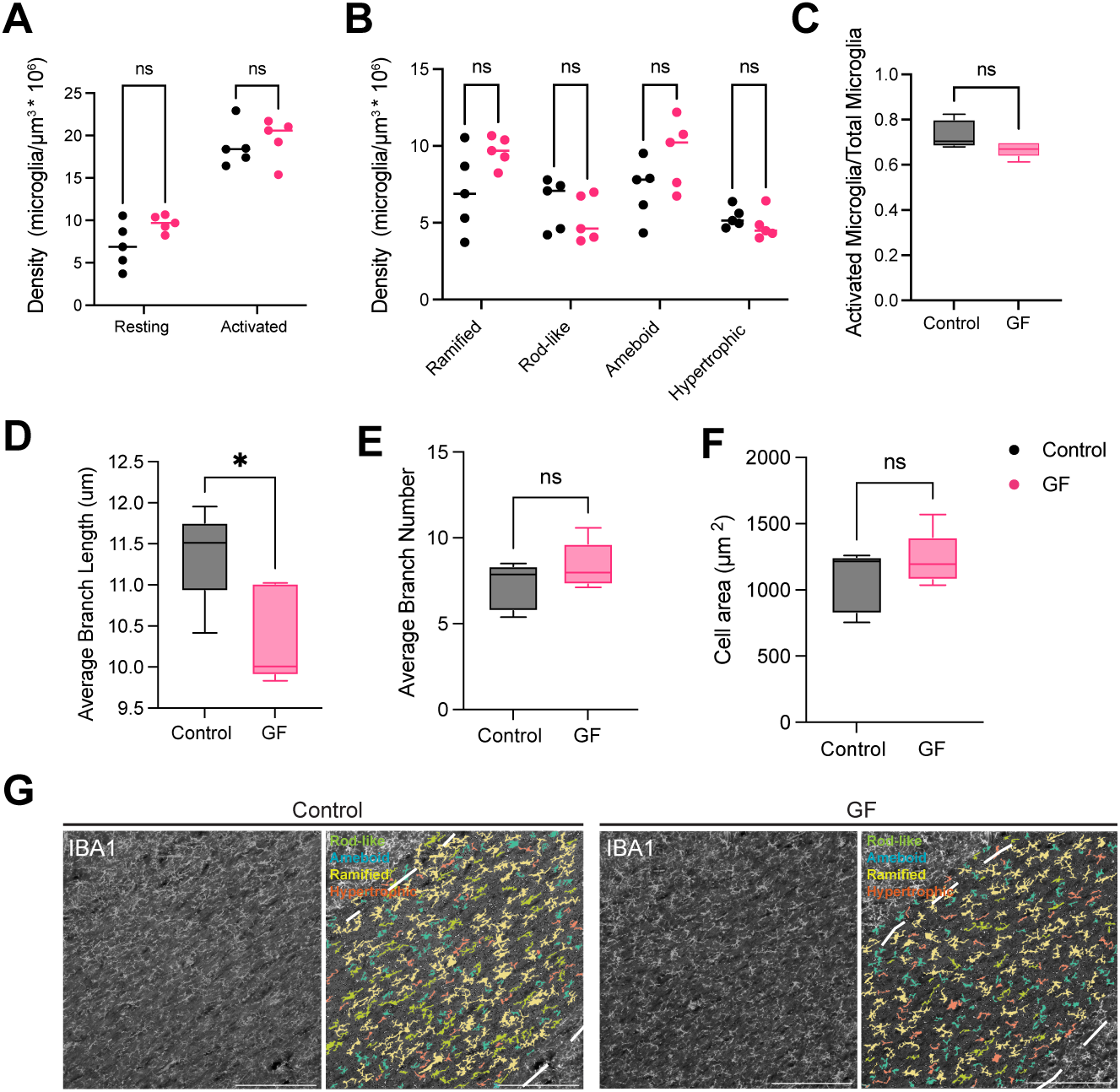
Microglia display a significant reduction in branch length in the PFCSWM under germ-free conditions. Analyses using microgial morphology data assessed by MicrogliaMorphology software program within the PFCSWM. **A,** Quantified density of activated and resting microglia between control (black) and GF (pink) tissue. **B,** Quantified control and GF microglial density within morphological groupings (ramified, rod-like, hypertrophic, ameboid). **C,** Proportion of activated microglia out of total microglia between control and GF tissue. Quantified average branch length (µm)**(D)**, branch number **(E)**, and cell area (µm^3^)**(F)** of individuals between Control and GF groups. **G,** Immunohistochemical images of the P16 PFCSWM in control (left 2 panels) and GF (right 2 panels) individuals are shown. Iba1^+^ staining (left) and individual microglia labeled by MicrogliaMorphology Colorbycluster macro (right) as rod-like (green), ameboid (blue), ramified (yellow), or hypertrophic (pink). White dashed lines indicate regional boundaries. Scale bars, 200µm. Density is calculated as microglia/µm^3^*10^6^. Error bars indicate mean ± SEM. ns *=* non-significant (*p* > 0.05), \**p* < 0.05.

### Microglia exhibit an activated morphology in the PFCII-III of germ-free piglets

Although the role microglia play in phagocytosing synapses has recently been challenged (O’Keeffe et al., 2025), several studies in rodents report significant disruptions in microglial synaptic pruning within the cortex of germ-free, antibiotic depleted, and mice with gut dysbiosis (Diaz Heijtz et al., 2011; Luck et al., 2020). These disruptions in synaptic pruning may contribute to the imbalances in excitation/inhibition commonly seen in disorders such as Autism Spectrum. In our previous GF study, we found no significant macrostructural differences in prefrontal cortical volume by MRI that may suggest massive loss of cortical neurons(Ahmed et al., 2021); however, we did not make any assessments at the cellular level that would reveal subtle differences underlying notable behavioral deficits in gut-brain axis animal paradigms. Since there are no reports on cortical microglia during perinatal development in GF piglets we aimed to determine if the activated/phagocytic phenotype documented in rodents is phylogenetically conserved in swine.

In the PFCII-III, GF animals displayed a higher density of microglial grouped as “activated” (activation state*□*×*□*treatment, *F*_(1,20)_□=□7.994, *p□=□*0.0104; activation state, *F*_(1,20)_□=□19.97, *p□=□*0.0002; treatment, *F*_(1,20)_□=□3.679, *p□=□*0.0695, two-way ANOVA) **(Fig. 5A),** as well as significantly higher densities of hypertrophic microglia (morphological state*□*×*□*treatment, *F*_(3,40)_□=□5.661, *p□=□*0.0025; morphological state, *F*_(3,40)_□=□132.2, *p□*<*□*0.000000000000001; treatment, *F*_(1,40)_□=□4.765, *p□=□*0.0350, two-way ANOVA) **(Fig. 5B),** and a larger proportion of microglia identified as morphologically activated (*t*_(10)_ *=* 2.496, *p* = 0.0317, unpaired *t* test) **(Fig. 5C),** indicating that animals in the GF group displayed a unique skew in the proportion of their microglia identified as activated. While there was no significant difference in average branch lengths (*t*_(10)_ *=* 0.5833, *p* = 0.5726, unpaired *t* test) **(Fig. 5D),** unpaired *t*-tests indicate that microglia in the GF tissue had significantly lower average number of branches and cell areas (*t*_(10)_ *=* 2.891, *p* = 0.0161, unpaired *t* test), (*t*_(10)_ *=* 3.563, *p* = 0.0052, unpaired *t* test) **(Fig. 5E, F).** These results indicate that the GF paradigm results both in increased densities of activated microglia as well as increases in individual features traditionally associated with activated and migratory morphology.

**Figure 5.**
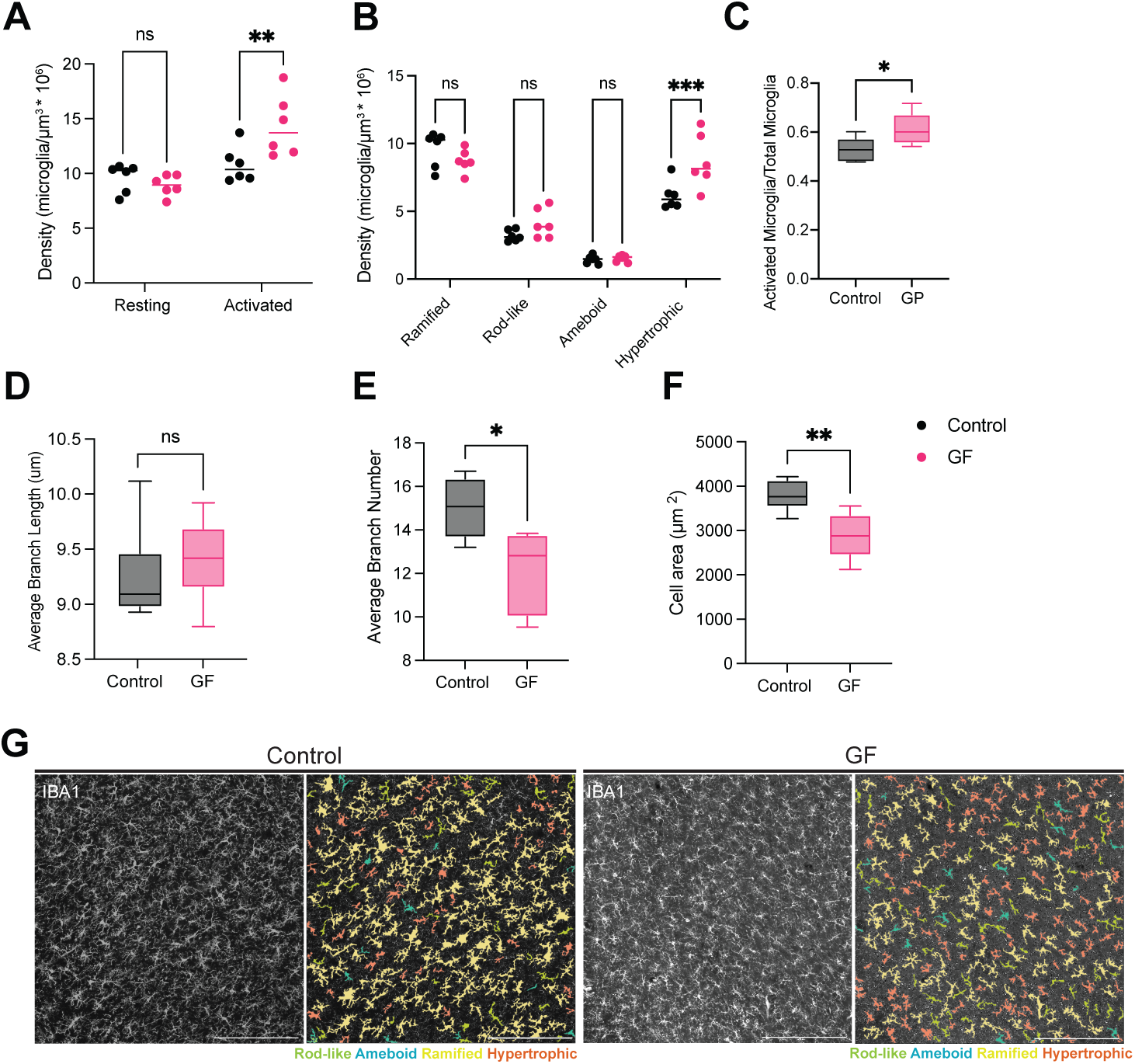
Microglia exhibit an activated morphology in the PFCII-III of germ-free piglets. Analyses using microgial morphology data assessed by MicrogliaMorphology software within the P16 PFCII-III. **A,** Quantified density of activated and resting microglia between control (black) and GF (pink) tissue. **B,** Quantified control and GF microglial density within morphological groupings (ramified, rod-like, hypertrophic, ameboid). **C,** Proportion of activated microglia out of total microglia between control and GF tissue. Quantified average branch length (µm)**(D)**, branch number **(E)**, and cell area (µm^3^)**(F)** of individuals between Control and GF groups. **G,** Immunohistochemical images of the P16 PFCII-III in control (left 2 panels) and GF (right 2 panels) individuals are shown. Iba1^+^ staining (left) and individual microglia labeled by MicrogliaMorphology Colorbycluster macro (right) as rod-like (green), ameboid (blue), ramified (yellow), or hypertrophic (pink). Scale bars, 200µm. Density is calculated as microglia/µm^3^*10^6^. Error bars indicate mean ± SEM. ns *=* non-significant (*p* > 0.05), \**p* < 0.05, \*\**p* < 0.01, \*\*\**p <* 0.001.

### GF paradigm results in upregulation of genes associated with an elevated immune response, microglial activation, and migration

Knowing that the PFC II-III appears to be uniquely vulnerable to changes associated with the GF paradigm, we next aimed to examine transcriptomic gene expression changes resulting from this. PFCII-III gray matter tissue was collected from animals and subsequently processed by immunoprecipitation. We then performed RNA-seq expression analyses to evaluate what, if any, transcriptomic changes could be observed within this tissue. This analysis revealed 37 differentially expressed genes (DEGs) that were annotated in the pig genome using *p* < 0.05 and a threshold of 1.2-fold change in expression as a cutoff for analysis **(Extended Data Table 6-1),** including *CD38* (p < 0.001, FC = 1.55), *RSAD2* (p < 0.001, FC = 0.08), *GABRR1* (p < 0.001, FC = 0.08), *FN1* (p < 0.001, FC = 0.54), *IFI6* (p < 0.001, FC = 0.37), *IRAK1BP1* (p < 0.001, FC = 1.74), *RGS1* (p < 0.001, FC = 3.22) **(Fig. 6A)**. Notably, one of these DEGs is *CD38* **(Fig. 6A)**, an ectoenzyme highly expressed in microglia that has been shown to increase in expression following induced neuroinflammation *in-vivo* (Roboon et al., 2021), as well as promote microglial activation and activation-induced cell death *in-vitro* (Mayo et al., 2008). Other notable genes such as *RGS1*, a microglia migratory gene (Atwood et al., 2011; Cunningham et al., 2013), and *IRAKBP1* which is required for TNF activation of NF-κB (Luo et al., 2007), were also upregulated. In turn, *FN1* or fibronectin 1 was significantly downregulated, which is not expressed in microglia, but has previously been showed to be engulfed by microglia during brain development (Lawrence et al., 2024) (**Fig.6A, Extended Data Table 6-1**).

**Figure 6.**
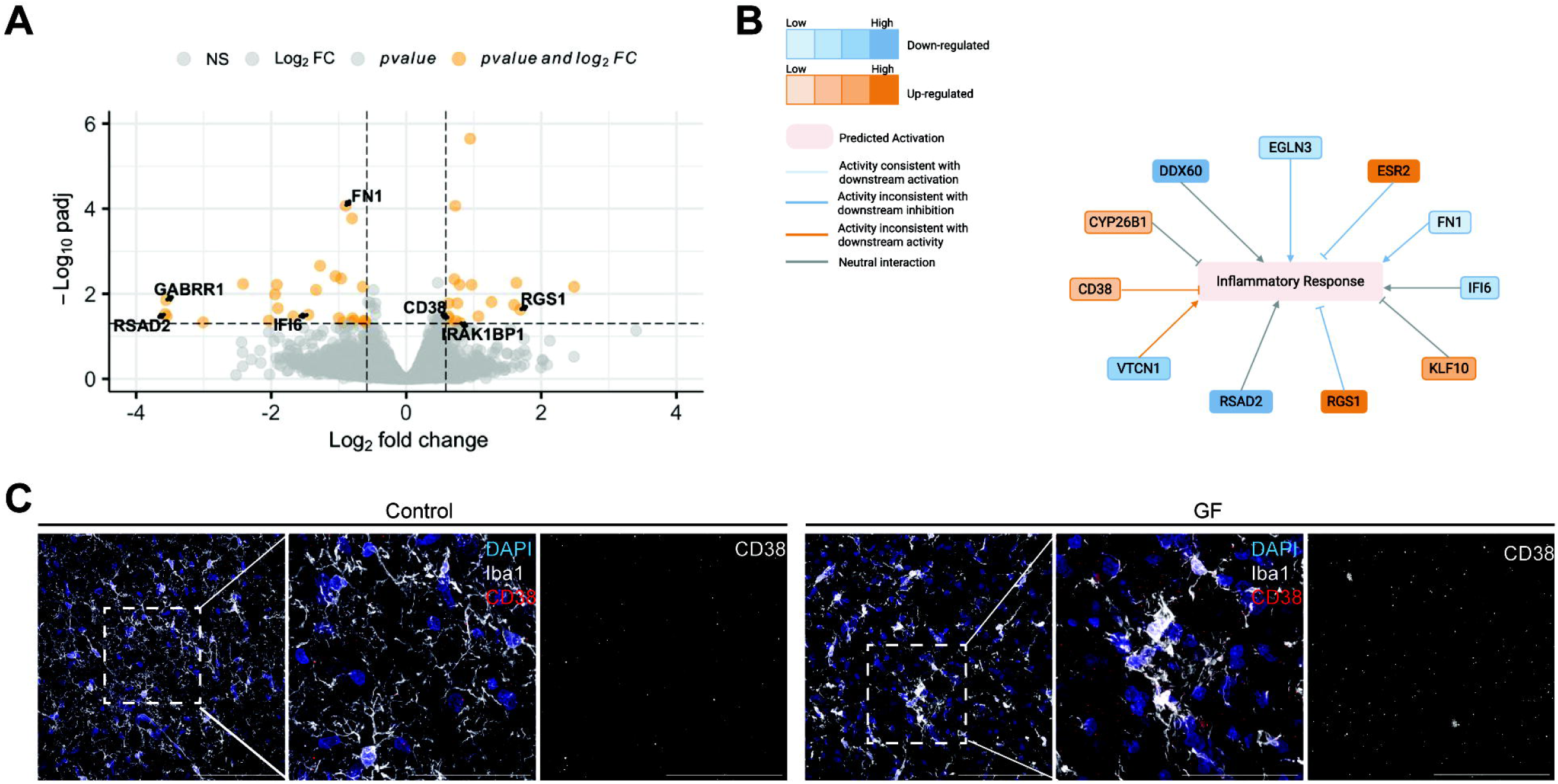
GF paradigm results in upregulation of genes associated with an elevated immune response, microglial activation, and migration. **(A)** Volcano plot of significant upregulated and downregulated genes in GF tissue, using p < 0.05 and a threshold of 1.2-fold change in expression as a cutoff for analysis. CD38 (p < 0.001, FC = 1.55), RSAD2 (p < 0.001, FC = 0.08), GABRR1 (p < 0.001, FC = 0.08), FN1 (p < 0.001, FC = 0.54), IFI6 (p < 0.001, FC = 0.37), IRAK1BP1 (p < 0.001, FC = 1.74), RGS1 (p < 0.001, FC = 3.22) are labeled in bold. **(B)** Graphical reconstruction of immune activation Ingenuity Pathway Analysis result, showing differentially expressed genes in our tissue related to this pathway. Panel made with BioRender. **(C)** Representative immunohistochemical images of the PFCII-III in control and GF animals. CD38^+^ (red), DAPI^+^ (blue), and Iba1^+^ (white) staining (left, scale bar 200µm), a middle inset of the leftmost image (middle, scale bar 50µm), and a single channel (CD38) of this inset (right, scale bar 50µm) for each group n=3/animals per group. Data supported by Extended Data Table 6-1.

Further examination with Ingenuity Pathway Analysis (IPA) showed that the top biological function of our dataset included immune activation. The analysis generated a graphical summary including the genes relevant to this function, including *CD38* **(Fig. 6B).** Performing IHC on control and GF tissue, we found increased CD38 staining on microglia in GF tissue in PFC (**Fig. 6C**). Taken together, these data support the inflammatory susceptibility of the PFC II-III, highlighting a profile of upregulated transcriptomic microglial/neuroinflammatory genes within GF tissue.

## Discussion

Microglia are known to coexist in multiple functional states throughout neurodevelopmental epochs and across different brain regions. Morphological features have been characterized to distinguish traditional functions such as resting/ramified and active/ameboid. However, there is a growing consensus that microglia are more functionally dynamic and require a combinatorial analysis of several morphological measures to fully capture their implied functional status *in situ* (Paolicelli et al., 2022). New efforts to develop automated pipelines capable of characterizing morphological states enable a standardized, unbiased, high-throughput approach to group microglial to infer functional states determined by open-access software. Because microglial functional states vary across species in a sexually dimorphic manner (Hanamsagar et al., 2017; Geirsdottir et al., 2019), we employed such a pipeline to preclude feature-selection bias and make apt comparisons to other well-studied species across neurodevelopmental paradigms.

In our study, we utilized MicrogliaMorphology (Kim et al., 2024) to evaluate microglia morphology in piglets within three key brain regions that are uniquely similar to the human equivalent during early toddlerhood. In addition, we challenged microglial homeostasis using a germ-free piglet model whereby an absence of microbiota is anticipated to result in an immature, more ramified phenotype as seen in rodents (Erny et al., 2015) and were able to parse out subtle and overt region-specific differences. Surprisingly, our transcriptomic signatures and morphological data suggest that PFC microglia in GF piglets were more ameboid and with increased inflammatory gene expression indicating differences in timing, neurodevelopment, and species may be important in assessing the impact of the gut microbiota on the CNS.

To date, few studies have evaluated microglia in swine at the cellular and transcriptomic levels, all of which were performed with different breeds (Geirsdottir et al., 2019; Sobierajski et al., 2022; Shih et al., 2023). A single wild pig and a group of male piglets (White x Landrace x Duroc cross) were used to generate cross-species single-cell microglial comparisons and across specific brain regions (Geirsdottir et al., 2019; Shih et al., 2023). Morphological assessments of microglia have been reported in the European Wild Boar spanning antenatal and postnatal neocortical development (Sobierajski et al., 2022); the closest comparable age assessed in this study was from one male boar at P30 where microglia were characterized in the SVZ on the level of the dorsoparietal cortex. Unlike the dorsoparietal cortex, which is nearly fully developed in swine at birth, the prefrontal cortex assessed here is less developed at birth and continues along a protracted timeline as new challenges arise during aging. In addition, our study was performed with a mix of domestic female and piglets (Landrace x Yorkshire) reared in SPF conditions and settings that do not resemble a wildlife setting. Therefore, it is difficult to make direct comparisons between our studies, especially considering the well-established sexually dimorphic nature of microglia heterogeneity and function in less commonly studied higher-order species.

It has become increasingly clear that microglial heterogeneity across species, sex, and context is difficult to compare between studies as nomenclature and quantitative approaches/measures vary widely. In the context of gut brain axes studies, increased ramification has been described as an immature phenotype in the mouse cortex as well as other brain regions assessed and more resistant to mounting an immune response after an LPS bacterial or viral challenge (Erny et al., 2015; Leclercq et al., 2017). While an ameboid morphology is often associated with phagocytosis (Nemes-Baran et al., 2020; Dayananda et al., 2023), it has been shown that ramified microglia can also phagocytose via their terminal branches during adult neurogenesis (Sierra et al., 2010). In addition, it is unclear whether a rod-like morphology overlaps with an activated phenotype versus a migratory phenotype. While our findings are suggestive of functional cellular activity, there are counter examples to consider.

Within the VZ/SVZ we found no significant differences in microglia numbers of morphology with a majority (∼80%) of microglia classified as activated suggesting no disruptions in ongoing stem cell pruning (**Fig. 3**). Within the PFCSWM we found no differences in microglia categorized as activated between groups; however, there was a significant reduction in branch length (**Fig. 4**) which has been linked to elevated cytokine release in postnatal rat hippocampal slice cultures (Madry et al., 2018). It has been documented that microglia display an ameboid morphology during phagocytosis of OPCs in the mouse corpus callosum (Irfan et al., 2022); therefore, it is unclear whether a reduction in branch length in the PFCSWM of the postnatal swine represents a reactive/responsive, less motile, or less mature state and whether this phenomenon is unrelated to our previous work demonstrating a reduction in OPC proliferation within this white matter tract (Ahmed et al., 2021). Future studies assessing microglial morphology paired with biochemical assays to corroborate activation status will be an informative step towards elucidating the impact of the microbiome on postnatal white matter brain development.

Here we report a significant shift towards an activated state in microglial within the GF PFC defined by an increase in hypertrophic morphology **(Fig. 5B),** a reduction in cell area **(Fig. 5E),** and a reduction in the average number of branches **(Fig. 5F).** Unlike the other regions assessed, there was an even proportion of microglia classified as activated (50%) to resting (50%) versus ∼80% active in the PFCSWM and VZ/SVZ **(Figs. 3C, 4C, 5C)** indicating that microglia may play more diverse roles in the PFC at an age when this cortical domain is undergoing interneuronal integration and synaptic refinement. Thus, a shift towards an activated state in piglets born and reared in germ-free conditions may represent a delay in microglial maturation -as the immune system isn’t primed by maternal microbes at birth- disrupting synaptic pruning capacity or hyperactive microglia phagocytosing excessive neurons or early immune responsive astrocytes. This regional difference in morphology has also been described in humans, where the PFC was described as less complex than other regions, regardless of sex (Yoblinski et al., 2025). Abnormal synaptic pruning and alterations in microglial morphology associated with function have been documented in the prefrontal cortex of dysbiotic mice whom where treated with acute, high doses of antibiotics in adulthood (Krishnapriya et al., 2025a; Krishnapriya et al., 2025b).

Our RNAseq findings revealed a significant elevation in differentially expressed inflammatory genes within the PFC of GF piglets in support of an immune activated microenvironment **(Fig. 6).** GF mice have been shown to have reductions in proinflammatory cytokine mRNA levels (Castillo-Ruiz et al., 2018). Future longitudinal studies employing single cell RNAseq on cell sorted microglia will be necessary to rule out confounding factors including blood barrier leakage, peripheral immune contributions, and intracellular vs. microenvironmental alterations in cytokine levels in GF animals. In addition, it will be important to corroborate whether elevated genes associated with an immune response play a classic inflammatory or an unknown role during brain development in the germ-free swine model.

Several studies in rodents demonstrate a strong link between gut dysbiosis and ASD-like behavioral phenotypes (Tochitani et al., 2016; Leclercq et al., 2017; Eltokhi et al., 2020). The piglet represents a phylogenetically tractable animal model to bridge comparisons between mice and humans in uniquely controlled experimental contexts. Our finding of a more activated phenotype in the PFC **(Figs. 5 & 6)** of germ-free piglets is in line with a spatial transcriptomic study showing elevations in reactive microglia within the cortex of ASD patients (Voineagu et al., 2011; Gupta et al., 2014; Parikshak et al., 2016; Gandal et al., 2022; Wamsley et al., 2024). We found no significant differences in cell area within the PFCWM in germ-free animals unlike the enlarged soma documented in microglia in the white matter of ASD cases (Morgan et al., 2010). Because ASD is predominantly a male disorder and postmortem assessments of human tissue are often at older ages, and our findings are derived from mixed sex cohorts, we make these comparisons with speculation. Future studies with a large number of male and female pigs to capture sexually dimorphic differences during childhood and through adolescence, paired with clinically relevant diagnostic measurements, will be essential in elucidating the underlying pathogenesis of neurodevelopmental diseases/disorders involving microglial flux.

## Extended Tables and Figuures

**Figure 2-1. Comparisons of microglial density between control and GF groups, split by region. (A-C)** Quantification of GF and control tissue microglial density within the VZ/SVZ **(A)**. Quantification of GF and control tissue microglial density within the PFCSWM **(B)**. Quantification of GF and control tissue microglial density within the PFCII-III **(C)**. Density is calculated as microglia/µm^3^*10^6^. Error bars indicate mean ± SEM. ns = non-significant (p > 0.05), *p < 0.05, **p < 0.01, ***p < 0.001, ****p < 0.0001.

**Table 6-1. Differentially expressed gene lists comparing GF to control tissue.** n = 3 per group.

## Abbreviations

ASD: autism spectrum disorder
CNS: central nervous system
GF: germ-free
MS: multiple sclerosis
NSPCs: neural stem progenitor cells
P: postnatal day
PFC: prefrontal cortex
PFCII-III: PFC cortical layers II + III
PFCSWM: PFC subcortical white matter
SVZ: subventricular zone
VZ: ventricular zone

## Author Contributions

BAL planned and performed experiments, analyzed data, and wrote the first draft and revised the manuscript. CK planned and performed experiments, analyzed data, and revised the manuscript. SNH, IPE, SEC, MRH, TP, TDA, JMB, and JPM performed experiments. AMP performed experiments, wrote, and revised the manuscript. PDM conceived the project, planned experiments, analyzed data, secured funding for the project, and wrote and revised the manuscript. All authors read and approved the manuscript.

## Acknowledgements

We thank the National Institutes of Health R01ES035013 and R15NS108183 (PDM) for supporting this work.

