## Supplementary figures and images for "Germ-free piglets display variable neuroinflammatory-like perturbations in prefrontal cortical microglia"

### Figure 2-1

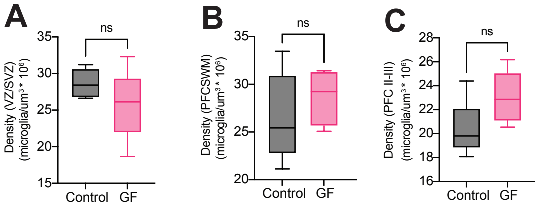
